# A novel computational approach to identify cancer cells in scRNA-seq data

**DOI:** 10.1101/2022.04.28.489880

**Authors:** William Gasper, Francesca Rossi, Matteo Ligorio, Dario Ghersi

## Abstract

Single-cell RNA-seq is an invaluable research tool that allows for the investigation of gene expression in heterogeneous cancer cell populations in ways that bulk RNA-seq cannot. However, normal (i.e., non tumor) cells in cancer samples have the potential to confound the downstream analysis of single-cell RNA-seq data. Several existing methods for identifying tumor cells use copy number variation inference. This work aims to extend existing approaches for identifying cancer cells in single-cell RNA-seq samples by incorporating putative driver alterations. We found that putative driver alterations can be detected in single-cell RNA-seq data and that a subset of cells in tumor samples are enriched in putative driver alterations as compared to normal cells. Furthermore, we show that the number of putative driver alterations and inferred copy number variation are not correlated in all samples. Taken together, our findings suggest that combining copy number variation inference with putative driver mutation load can augment the number of tumor cells that can be confidently included in downstream analyses of single-cell RNA-seq datasets.

## Introduction

Single-cell transcriptomics is revolutionizing the way we look at cancer. The ability to measure gene expression at the single cell level is particularly important given the high degree of intratumor heterogeneity exhibited by human cancers [1]. Intratumor heterogeneity – often promoted by genomic instability – leads to the emergence of different cancer cell subclones, some of them promoting metastasis, drug resistance, and disease recurrence [2]. The molecular characterization of cancer subclones via single-cell transcriptomics is therefore essential for achieving more effective, personalized treatment options and for improving clinical outcomes.

A fundamental prerequisite for an effective downstream analysis of single-cell transcriptomics datasets is the correct identification of cancer cells, which is often complicated by low tumor purity (i.e., low proportion of tumor cells in cancer samples). In addition to low tumor purity, the cellular process known as epithelial-to-mesenchymal transition (EMT) can further complicate the identification of tumor cells [3]. EMT may cause epithelial cells to express fibroblast-like transcriptional programs and may make (*i*) cancer-associated fibroblasts virtually indistinguishable from cancer cells undergoing EMT, or (*ii*) non-EMT cancer cells indistinguishable from the normal epithelial cells in the surrounding organs (e.g., breast, prostate, pancreas) [3].

Bioinformatics methods are frequently used to selectively call cancer cells in single-cell transcriptomics datasets. Some of these methods rely on the analysis of marker gene expression to select normal cell types that can be confidently identified [4] or perform gene expression-based clustering [5]. While these methods add valuable information to the cell filtering process, they leave the potential for both false positives and false negatives and may not be useful for identifying cancer cells in epithelial and other cell populations. In addition, the EMT phenomenon mentioned above can reduce the effectiveness of gene expression-based approaches.

Other *in silico* tools infer tumor cell-specific genetic variants, such as copy number variation (CNV), from scRNA-seq data. CNV is defined by duplications or deletions of chromosome parts and is considered a cancer-specific signature often associated with clinical outcomes [6]. One example of a popular CNV-based tool is InferCNV, which was developed to infer large-scale CNV for glioblastoma cells in Smart-Seq generated transcriptomics data [7]. Other tools include HoneyBADGER [8], CaSpER [9], and CopyKAT, which identifies CNVs at a resolution of 5 Mb and is suitable for 10X Genomics-generated scRNA-seq data [10].

CNV-based tools have proven effective at calling cancer cells for a variety of malignancies. However, CNV-inference-based filtering has limited efficacy when applied to samples from certain cancers commonly characterized by limited CNV. Examples of low-CNV cancers include prostate cancer [11], thyroid carcinoma, and clear cell renal carcinomas [12]. Even in tumor types commonly characterized by medium-to-high CNV, these methods would falsely exclude tumor subclones with low CNV, which we have identified in this work.

To address these issues, we propose a novel computational framework for identifying cancer cells in transcriptomics data. Our approach focuses on inferring two distinct classes of genomic events that are largely associated with tumorigenesis and tumor progression: driver single nucleotide variants (SNVs) and copy number variation (Figure 1). By adding SNVs, our method is particularly useful when attempting to identify cancer cells in low CNV tumors and for the identification of cancer cells with potentially targetable driver mutations, in both low and high CNV tumors. Our presented methodology and findings suggest that analysis of variants identified in cancer scRNA-seq data is underutilized and may yield novel insights and applications.

**Fig 1.**
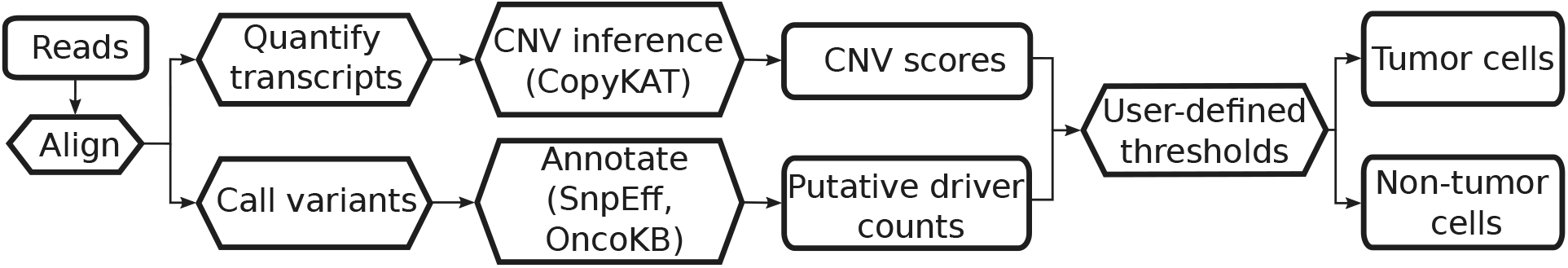
Flowchart depicting the general methodology described in this work. Rectangles indicate inputs and outputs, hexagons indicate processes.

## Materials and methods

### Data selection

Smart-seq2-generated scRNA-seq data from triple negative breast cancer (TNBC) samples and corresponding whole-exome sequencing (WES) results were obtained from the Gene Expression Omnibus (accession `GSE118390` [13]) As a basis for comparison, healthy pancreas, liver, dermal fibroblast, and neutrophil cells, from samples prepared using Smart-Seq2, were retrieved from [14], [15], [16], and [17], respectively. 200 cells were randomly selected from each dataset, with the exception of the dermal fibroblast dataset, which contained only 190 cells. In order to evaluate the potential use of this method with data generated using 10X Chromium (v3.1 chemistry) library preparation, a dataset containing a 20,000 cell mixture of non-small cell lung cancer dissociated tumor cells was obtained from the 10X Genomics website (https://10xgenomics.com/resources/datasets).

### Data processing

#### Quality control

TNBC cells were initially filtered according to the methodology detailed in [13]: cells were selected based on library size, number of expressed genes, and total mRNA. For any of the prior metrics, cells were excluded if values fell below 4 median absolute deviations from the median. In addition to the methodology described in [13], the threshold for the minimum number of expressed genes per cell was raised to a constant value of 1,000, and an additional cell was excluded by CopyKAT [10] due to insufficient chromosomal coverage (at one or more chromosomes) to reliably infer CNV. This resulted in 942 viable cells for analysis. For variant calling, FASTQs were retrieved using the SRA Toolkit and Trim Galore [18] was used to remove adapters and low-quality basecalls from read ends. For normal tissue cells, 173 cells were excluded by CopyKAT due to insufficient chromosomal coverage, leaving a total of 617 normal tissue cells for normal cell CNV inference analysis. All 790 randomly-selected normal tissue cells were used to create a generalizable alteration counts distribution, in an effort to incorporate as much dataset heterogeneity (particularly, with respect to wet-lab variability and batch effects) as possible.

#### scRNA-seq processing

Reads were aligned to the most recent hg38 reference genome build, retrieved from UCSC, using STAR aligner [19]. BCFTools [20] was used to generate genome pileups and to call and filter variants with a minimum quality score of 30. Variants were then annotated using SnpEff [21]. After SnpEff annotation, coding sequence variants were annotated using the OncoKB API [22] to obtain any known or predicted oncogenicity for the variant. The OncoKB API use requires an API token, obtainable at https://oncokb.org/apiAccess. A complete pipeline producing these results is available in the project code repository.

#### Coverage quantification

Due to normal variations in gene expression, random dropout, capture inefficiencies, and other sources of noise, it cannot be assumed that SNVs are not present in a sample unless the sample has sufficient coverage at the corresponding genomic position. To this end, a custom Python script was used to determine whether it is possible to effectively call an SNV at any given genomic position, based on a required minimum read depth of 5 at the respective position. This minimum depth is only used to identify the absence of a variant (for Figure 3), as no assumptions about genome coverage can be made for transcriptomics data. For calls about variant presence, a minimum quality score of 30 was used, as described in the prior section.

After alignment and annotation, all putative driver alterations were extracted from annotated VCF files. For each residue-level putative driver alteration, the genomic positions corresponding to the residue were then looked up using the most recent hg38 GTF file (retrieved from UCSC) and stored. Then, for each sample, genome pileups were generated from sorted BAMs using BCFTools and piped into a Python script, which measured coverage at all relevant genomic positions. In analysis, this coverage data was used to determine whether a putative driver alteration was genuinely absent or if there was insufficient coverage to call either way.

### CNV inference

CNV inference was performed using CopyKAT [10] with default parameters and a window size of 100. The TPM matrix originally produced in [13] (available under `GSE118390`) was used as input for CopyKAT. For normal tissue, transcript quantification was performed using `htseq-count` from HTSeq [23] in intersection-strict mode, converted to TPM, and then CNV was inferred using CopyKAT.

### WES data processing

Whole-exome sequencing (WES) samples from TNBC patients used for validation were initially trimmed to remove adapters and low-quality basecalls using Trim Galore [18]. Reads were aligned to the most recent hg38 reference genome build using BWA [24] and sorted using SAMTools [20]. PCR duplicates were removed using GATK’s MarkDuplicates [25]. BCFTools was used for generating genome pileups, variant calling, and variant filtering. SnpEff and the OncoKB API were used for annotating final variant calls.

### 10X data processing

The 10X NSCLC 20k mixture dataset was initially processed using Cell Ranger [26] from 10X Genomics. For each cell, reads corresponding to each unique cell barcode were extracted from all sequencing runs and sorted. Variant calling proceeded similarly to the TNBC dataset processing: BCFTools was used to generate pileups and call variants, and variants were annotated using SnpEff.

## Results

### CNV inference

We performed CNV inference using CopyKAT [10] to assess the viability of cancer cell calling based solely on CNV inference. CopyKAT identified structural genomic heterogeneity both within and between patients, and the results appear similar to those generated using a custom methodology and presented in Karaayvaz et al. [13]. A significant number of cells had limited or no CNV, and CopyKAT predicted 135 out of 942 cells as aneuploid. For the cells predicted as aneuploid, all but one (belonging to PT081) belonged to PT039. These predicted aneuploid cells are clearly evident in Figure 2 as the cluster of PT039 cells with comparatively extreme expression scores.

**Fig 2.**
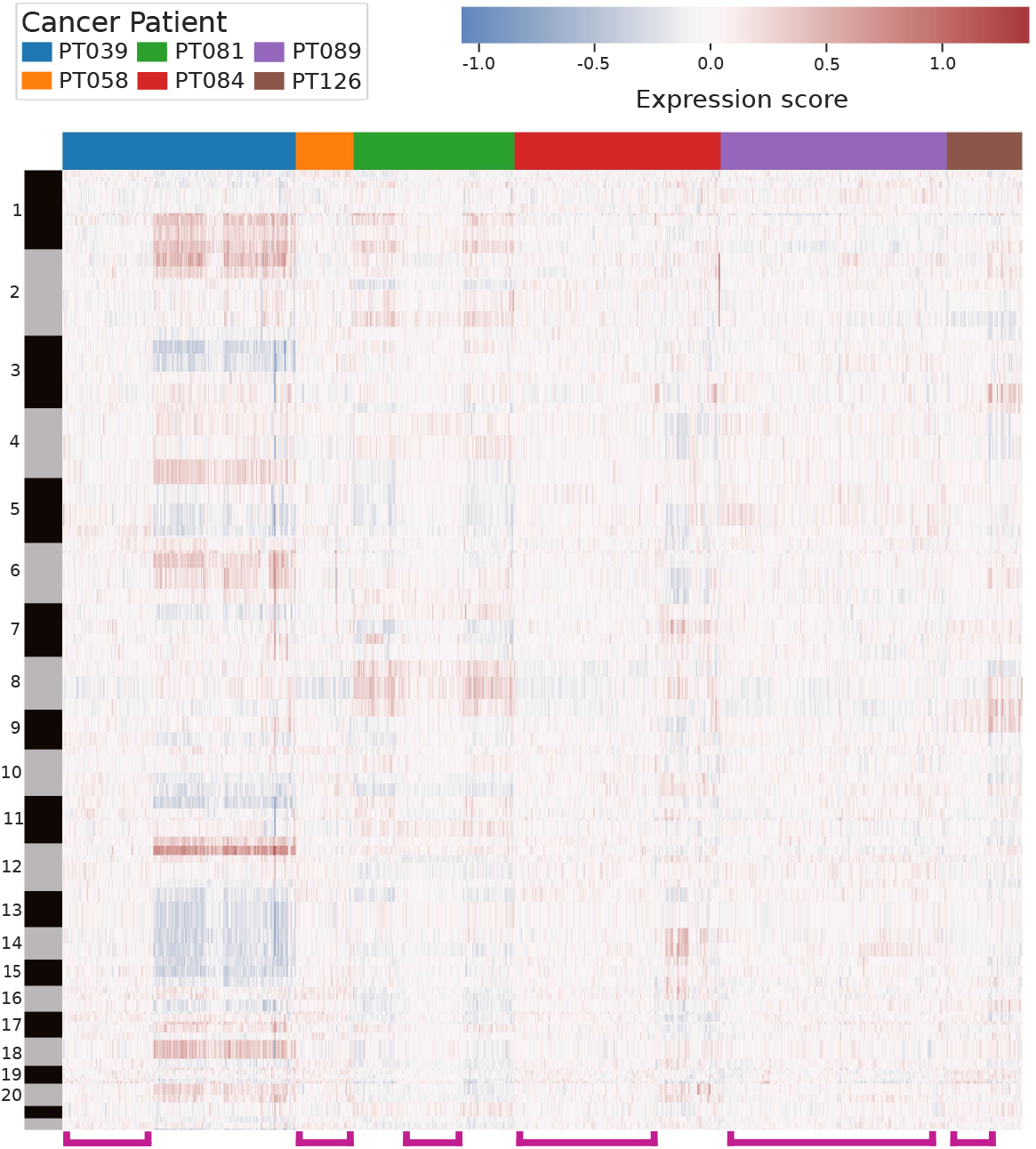
Heatmap visualizing inferred CNV results obtained using CopyKAT [10]. Columns represent individual cells, rows represent genes arranged by chromosomal position. Alternating bars on the y-axis indicate chromosome. Cells are clustered hierarchically by patient using expression score matrices. Pink brackets indicate cells of interest: cells with low inferred CNV that may be excluded during filtering.

**Fig 3.**
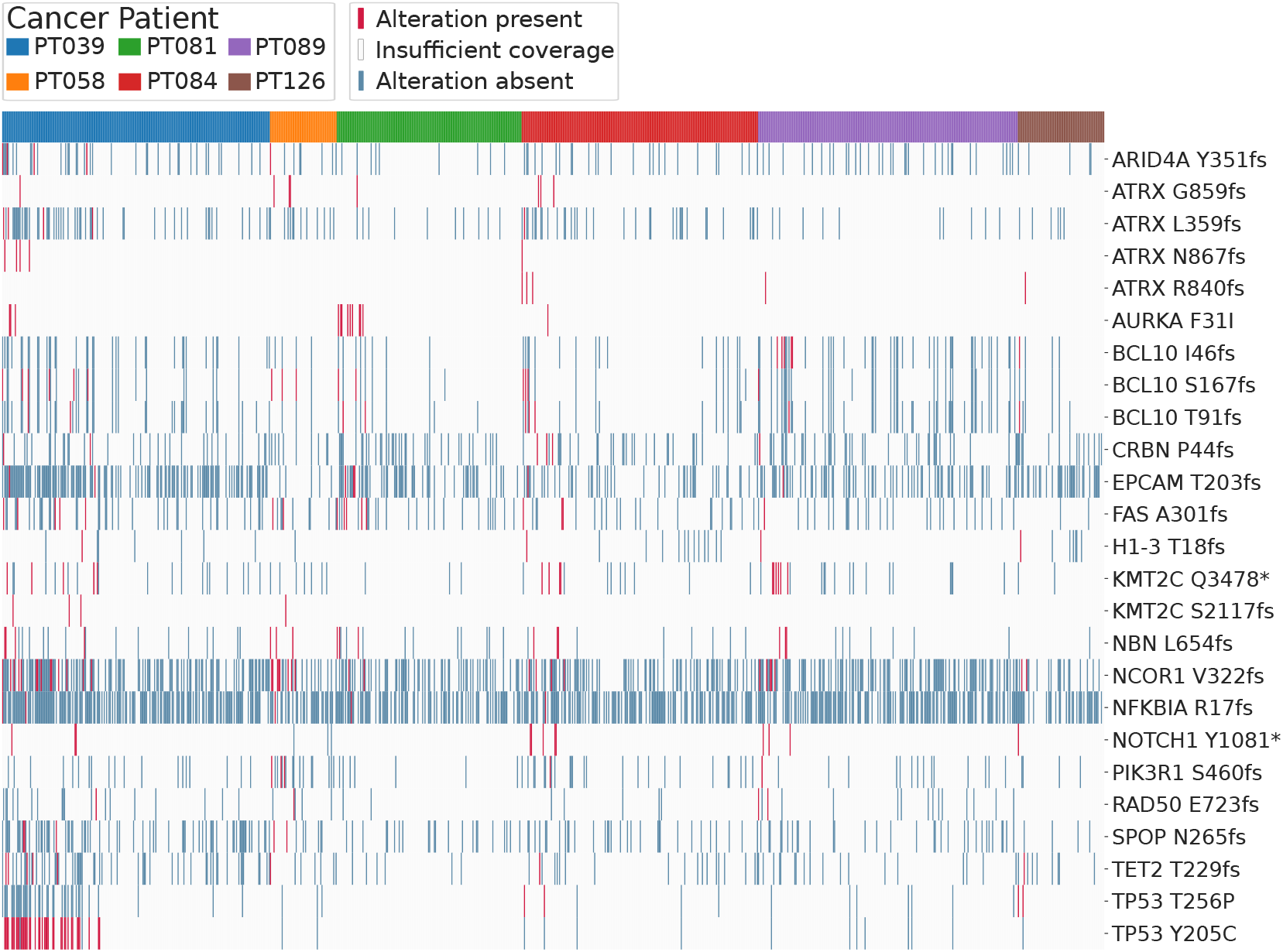
Heatmap indicating alteration status for 942 cells for the top 25 most frequent oncogenic, predicted oncogenic, and likely oncogenic alterations. Alterations are annotated using OncoKB. Absence of an alteration is noted when a cell has a read depth of at least 5 for all bases corresponding to the residue. For residues without an oncogenic alteration and with read depths less than 5 for all corresponding bases, the presence or absence of an alteration is not characterized (“Insufficient coverage”).

The presence of large, contiguous regions of high or low expression scores may indicate major structural alterations: either aneuploidy or considerable copy number alteration. Groups of cells predicted to have limited or no structural alterations are highlighted by pink brackets along the x-axis of Figure 2. Interestingly, all cells for patient PT058 had low CNV scores (Figure 2), with a mean of 0.08, and would likely be filtered out based on the CNV criterion alone. These results provide the rationale for including additional criteria like the presence of putative driver mutations.

### Variant calling and annotation

In order to investigate the possibility that cells predicted to have limited structural alterations are indeed cancer cells, we examined whether they harbored putative driver alterations. Variant calling on the TNBC data resulted in the identification of a number of putative driver alterations, defined as those alterations annotated by OncoKB as “Oncogenic", “Likely Oncogenic", or “Predicted Oncogenic". These results are visualized for the 25 most frequent putative driver alterations in Figure 3.

There are several examples of characteristic mutations common to multiple cells: for PT039, the TP53 Y205C (found in 41 cells), TP53 P72R (27), and NCOR1 V322fs (14) alterations; for PT081, the AURKA F31I (9) mutation; and for PT089, the KMT2C Q3478* (6). PT039 had a number of cells (24) containing multiple TP53 mutations, which seems to be a characteristic signature for these cells within this patient’s sample. There were also 122 alterations common to more than one and less than six cells per patient, and then a large number (1,467, non-unique) of singleton mutations occurring in only one cell per patient. These rare mutations are called based on high quality scores (*≥*30) and may represent genuine genomic variation, but should not be expected to be confirmed in WES or whole-genome sequencing, both of which capture genomic variation common to large numbers of cells.

A number of cells harbored no putative driver alterations. For all cells, a total of 1,454 unique putative driver alterations were identified with a median of 1.0 per cell. At least one SNV was identified in every cell, with a median of 2,897 SNVs per cell.

These results indicate that variant calling on scRNA-seq data with sufficient coverage can add complementary information into the cell filtering process. Cells with limited or no predicted structural alterations may be included based on predicted driver alterations. For example, PT058’s cells contain limited inferred CNV (Figure 2 & Supplementary Figure 7) but frequently contain predicted driver alterations (Figure 3 & Supplementary Figure 6). Based on these findings, we believe that actionable information can be added through variant calling and annotation. The addition of this data can allow for the inclusion of additional true cancer cells that might otherwise be excluded.

### Normal tissue analysis

We also investigated the presence of predicted oncogenic driver alterations in scRNA-seq data generated from normal, healthy patients for neutrophils (obtained from saliva), pancreas, dermal fibroblast, and liver cells in order to ensure that our putative driver alteration counts could be used to reliably identify cancer cells. Limited numbers of putative driver alterations were found in normal tissue scRNA-seq data: a plurality of healthy cells (52.3%) contained 0 putative oncogenic alterations, with an absolute maximum of 8 putative driver alterations, a mean of 0.89, and a median of 0.0. Putative driver alteration counts were found to be significantly different for TNBC and normal cells (Mann-Whitney U, *p* = 0.0023). Box plots comparing the normal cell putative driver alteration counts to TNBC patients are shown in Supplementary Figure 6. For CNV inference, the mean CNV expression score for normal tissue cells was 0.06 with an absolute maximum of 0.11. In comparison, the mean for the TNBC dataset was 0.09, and the distributions were found to be significantly different (Mann-Whitney U, *p* = 2.20 *·* 10^*−*84^). Box plots visualizing these distributions are shown in Supplementary Figure 7. Based on the prior results, we do not expect that non-tumor cells should contain the high numbers of predicted driver alterations or the high mean absolute CNV expression scores present in a number of the TNBC dataset cells.

### Relationship to inferred CNV

Having established that putative driver alterations are very rare or absent in normal cells, we set out to determine whether including putative driver alterations can effectively augment the number of predicted cancer cells when compared to CNV inference alone. In order to do this, we studied the relationship between putative driver alterations and inferred CNV in tumor samples (Figure 4).

**Fig 4.**
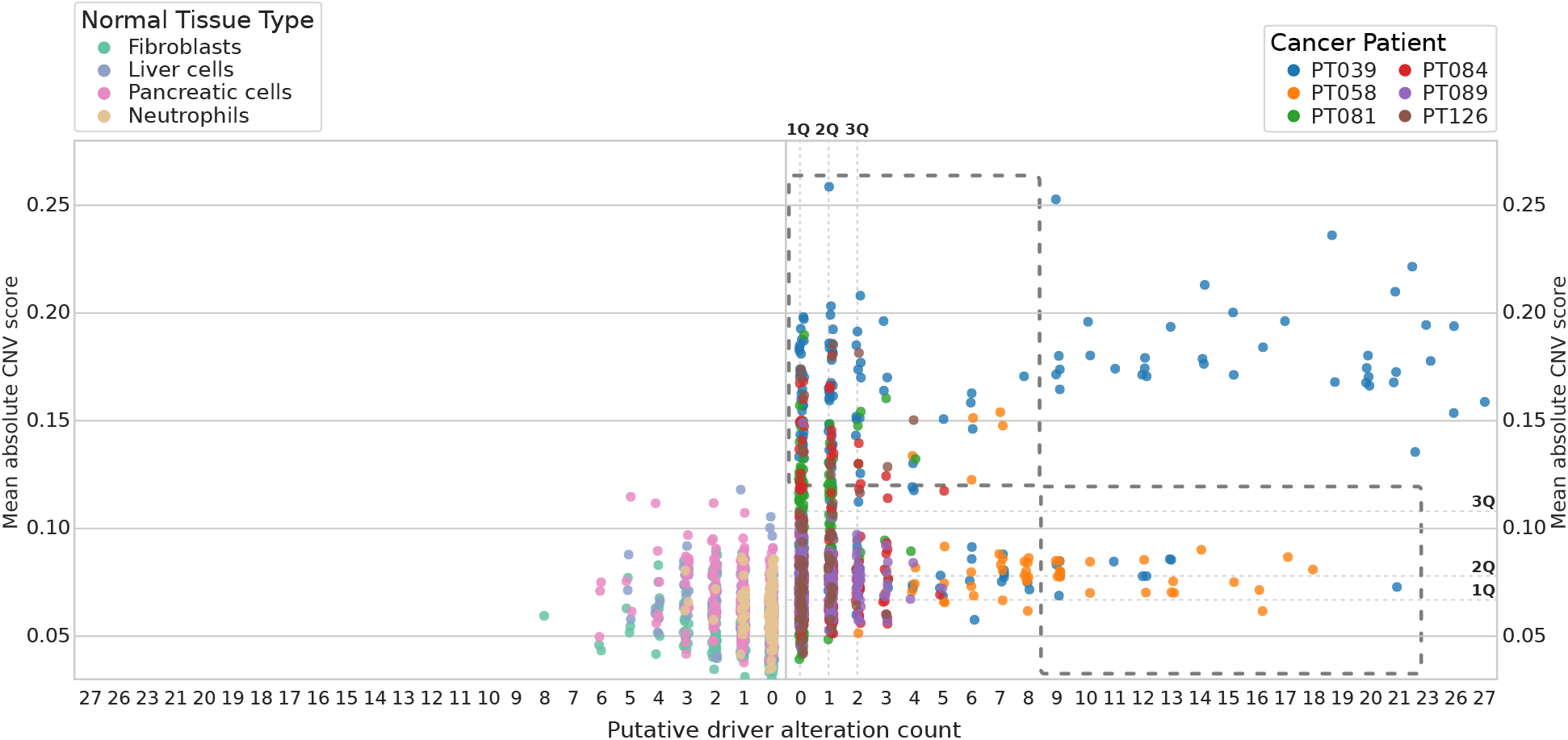
Relationship between putative driver alteration counts and inferred CNV for normal tissue (left) and tumor (right) dataset cells. Higher mean absolute CNV values indicate predicted structural alterations resulting in copy number variation, and lower values suggest limited CNV. Dashed rectangles indicate regions of interest: groups of cells that might be identified as cancer cells by either CNV inference or putative driver alteration count. First, second, and third quartiles are indicated for tumor cells by dashed lines and bold labels along their respective axes.

A number of cells with limited inferred CNV had high numbers of putative driver alterations. For example, 58 cells from the TNBC dataset are below the 99th percentile of normal tissue mean absolute CNV scores but above the 99th percentile of normal tissue putative driver alteration counts. Some of these cells of interest are indicated in Figure 4 by the lower dashed rectangle. These cells represent cells that might be excluded by CNV-inference-based filtering (and are predicted diploid by CopyKAT) but are likely to be tumor cells. For PT058, these predicted low-CNV cells represent the bulk of the data. Additionally, some cells from PT039 had limited predicted CNV but high numbers of predicted driver alterations. These cells may represent a distinct clonotype, an observation also made in [13] and supported by our CNV inference findings shown in Figure 2.

Overall, there appears to be low correlation between the putative driver alteration counts and mean absolute CNV expression score. However, this varies widely based on patient: PT058, PT084, and PT089’s cells had no correlation between CNV and alterations, and the relationship is weak, albeit significant, for PT081 (Table 1). This variability may be attributed to inter-patient somatic genomic heterogeneity.

**Table 1.**
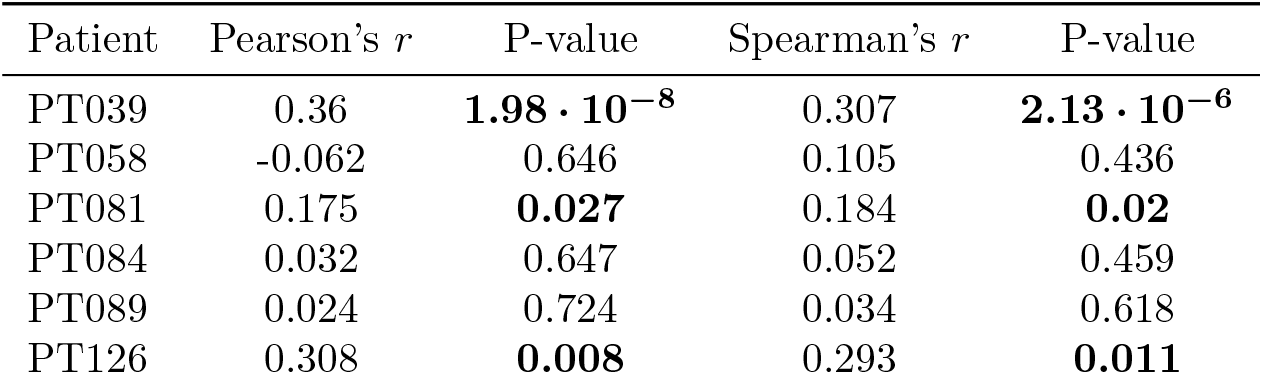
Table showing the correlation coefficients and statistical significance for relationships between inferred CNV and alteration counts for cells belonging to each patient. P-values less than 0.05 are shown in bold.

In the context of CNV inference, the addition of putative driver alteration counts represents an additional dimension that can be effectively used to separate cells. Cells that have similar predicted degrees of structural alteration may be on opposite ends of the alteration counts distribution, and this increased dimensionality allows for the separation of cells that appear similar solely in the context of inferred CNV.

### Comparison to WES

In order to further validate our findings, we performed variant calling and annotation on available WES data corresponding to a subset of TNBC patients. Processing and analysis of WES data resulted in fewer putative driver alterations than were found in the scRNA-seq data. However, a number of marker alterations common to cells for PT039 and PT081 were confirmed by the WES analysis, shown in Figure 5. For the putative driver alterations identified in WES, most were either found in cells from the corresponding patient or lacked sufficient coverage in most of the patient’s cells to definitively call presence or absence.

**Fig 5.**
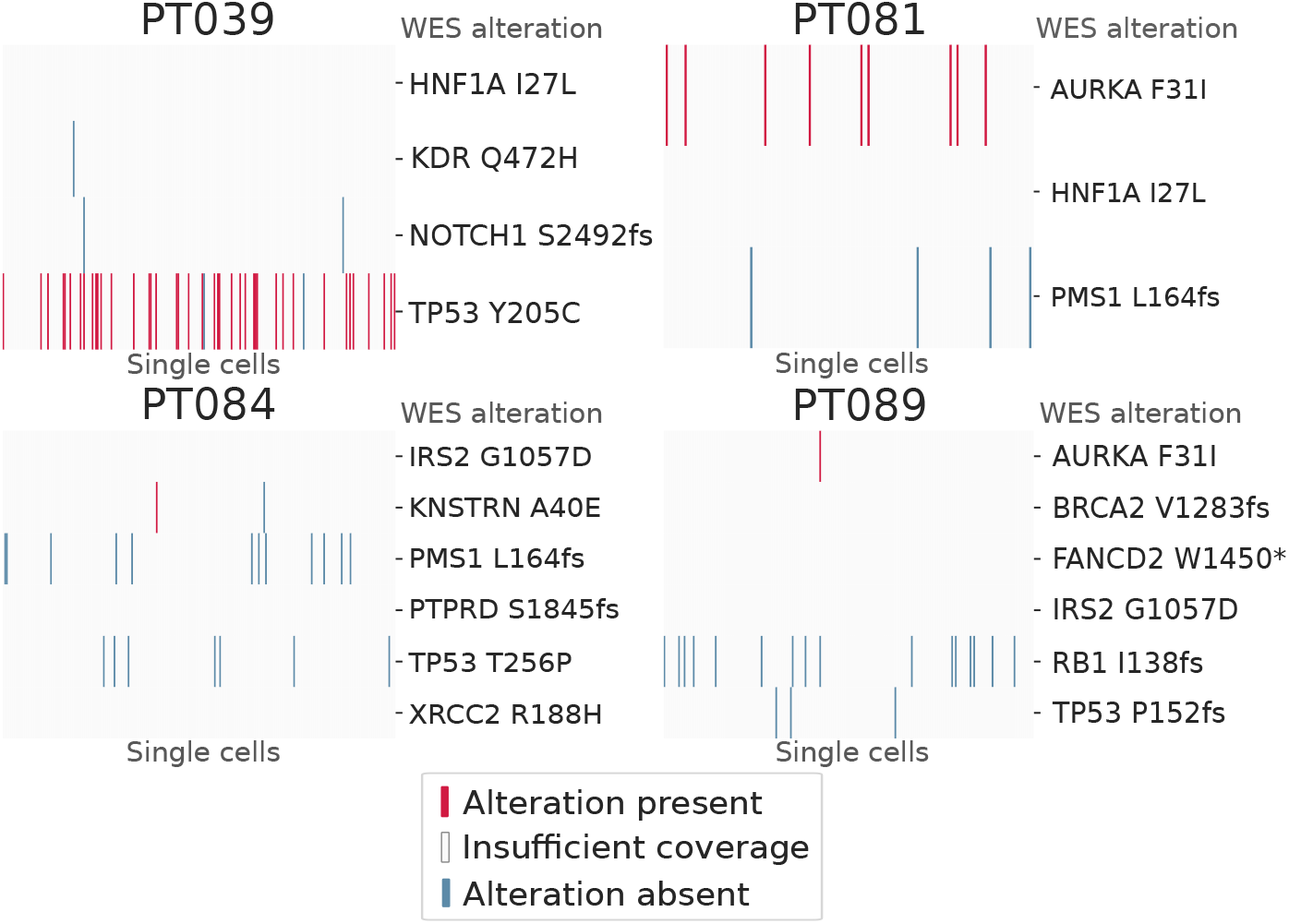
Heatmaps showing the overlap between putative driver alterations found in single cells and each patient’s respective WES samples.

There appear to be a few possible conflicts between the scRNA-seq and WES variants. For example, the PMS1 L164fs mutation found in PT084’s WES data does not appear in any of the patient’s cells, despite having sufficient coverage in some cells. Similarly, the RB1 I138fs mutation found in PT089’s WES data was not found in any cells, despite having coverage in many cells. These discrepancies may be due to intra-tumor genomic heterogeneity. Both our CNV inference results and those originally presented in [13] suggest that there may be multiple clonotypes for multiple patients, including PT084. Additionally, findings comparing the scRNA-seq-derived inferred CNV to the WES-derived inferred CNV (presented in [13]) suggest that clonotypes present in the scRNA-seq samples may not be presented in the WES. In general, the WES results support our findings in the scRNA-seq data: WES alterations were either found in a number of cells from the same patient or there was insufficient coverage to make the call.

### 10X Genomics NSCLC DTC dataset

10X Genomics scRNA-seq library preparation protocols are widely used and thus present an attractive target for this methodology. However, significantly fewer oncogenic alterations were found in the 10X Genomics NSCLC dataset. Only 16 putative driver alterations were found in reads corresponding to 13,093 unique cellular barcodes passing filtering in all sequencing runs. Our current methodology has limited efficacy when applied to this data, and likely does not provide an enriching effect with which to augment CNV inference on 10X-generated data.

## Discussion and Conclusion

This work highlights the viability of calling the presence and absence of oncogenic SNVs in scRNA-seq data. SNVs can be readily identified in scRNA-seq data generated by the Smart-seq2 protocol, and these SNVs may be used to increase confidence when filtering data to exclude non-tumor cells. This approach is particularly useful when cells lack major structural alterations (Figure 4 and Supplementary Figure 7). Although CNV inference will not likely yield false positives when selecting cells with a high score, it may result in false negatives.

Aside from CNV inference, other methodologies for cancer cell filtering may produce both false negatives and false positives. Cluster-based filtering may require arbitrary boundary decisions that may include or exclude cells incorrectly, and clonotypes represented by small numbers of cells may not produce distinct clusters. Marker gene expression may effectively allow for the exclusion of certain cell types, particularly immune and endothelial cells, but will not help to discriminate between tumor and normal epithelial cells. Additionally, tumor-macrophage fusion cells [27, 28] may be incorrectly called immune cells when using marker-based exclusion methods.

The results presented here illustrate the need for augmenting CNV inference with additional evidence to confidently call cancer cells in scRNA-Seq datasets. Our results showed that several cells with low inferred CNV harbored a number of oncogenic alterations, as shown in Figure 4. Additionally, there is not always a correlation between inferred CNV and the number of putative driver alterations identified (Table 1). If cell filtering on this data were based on CNV inference alone, then a number of likely-cancer cells would be excluded. This phenomenon is particularly pronounced in PT058’s cells, which had comparatively limited inferred CNV but frequently had numerous putative oncogenic alterations. We suspect that a number of these cells are genuinely cancer cells with limited CNV. Structural alterations are not requisite for tumorigenesis and disease progression, and tumors with minimal structural alterations but comparatively high mutation burden have been observed in prior work [29–31]. Due to the high data quality and quality control procedures, we believe that the CNV inference results are accurate, so these findings are not likely to be the result of an inability to effectively detect CNV.

Our work provides a way to bolster cell filtering with additional evidence. In isolation, this method may not be sufficient to identify cancer cells in a highly reliable way: normal expression variation and other sources of noise suggest that oncogenic alterations will not always be consistently present in the scRNA-seq data for cancer cells. However, our method effectively augments calls made using alternate methods and is especially interesting in the context of CNV inference data. The added dimension provided by putative driver SNV counts can effectively separate cells that are indistinguishable by CNV alone.

Unfortunately, there are a handful of current issues preventing the extensive use of this method with 10X-generated data. The 10X Genomics 3’ protocol generates reads that only cover a short region of the 3’ end of each transcript captured. This poses a serious limitation for the identification of oncogenic coding sequence alterations, which are the focus of this work. The majority of the target genes examined here have a 3’ untranslated region longer than the short reads typically sequenced in scRNA-seq experiments. However, there does seem to be potential for future studies in this area, as oncogenic alterations were indeed identified in a small proportion of cells in 10X Genomics data.

The presence of putative driver alterations in some normal cells is an interesting finding. While there may be comparatively less confidence in scRNA-seq variant calling compared to WES or WGS, we suspect that at least some of the variant calls represent genuine somatic mosaicism. There appears to be only a limited body of work quantifying the expected rates of short variant single-cell somatic mosaicism. Lodato et al. found varying rates of single-cell somatic SNV accumulation in neurons (up to *≈* 40 per year) and that single-cell SNV counts strongly correlated with age [32]. Large scale single-cell somatic mosaicism has also been found in neurons: McConnell et al. found that 13*−*41% of frontal cortex neurons harbor megabase-scale *de novo* CNVs [33]. Expected short variant single-cell somatic mosaicism rates for other normal tissues do not appear to be well characterized. Widespread single-cell short variant somatic mosaicism might be expected based solely on replication error rates [34,35], but there does not appear to be extensive experimental evidence to support this. These results indicate the need for additional experimental investigation into the normal cell space to confirm whether the SNVs found in data from healthy individuals (Figure 4 and Supplementary Figure 6) are due to PCR artifacts during library preparation or whether they represent true genetic alterations with potential unexplored biological implication.

Certain cancer types may be more amenable to our proposed methodology than others, and cancers with high mutation burdens may be the most viable targets. A number of cancer types, such as skin, lung, certain lymphomas, and various squamous cell carcinomas have comparatively high mutation burdens [36] and may be most likely to harbor large numbers of putative driver alterations. Additionally, characteristic, high-frequency mutations may be identified in cells to increase confidence in cancer status. These characteristic mutations may be limited to the context of specific patients, like the TP53 Y205C mutation in PT039’s cells (Figure 3), but may more broadly be found in certain cancer types: for example, KRAS G12x mutations can be found in 64.9% of pancreatic adenocarcinoma samples aggregated on CBioPortal [37].

In the future, we plan to leverage genomic inferences made from scRNA-seq variant calling in additional ways in order to increase the efficacy of this methodology. We also intend to provide a production-ready solution for end users to incorporate this methodology into analysis pipelines. The current methodology requires an OncoKB API key, and users wishing to adopt our methodology or reproduce our results will need to apply and obtain a token. Currently, we provide scripts to reproduce results presented here and a script that can be used to parse SnpEff output, count putative driver alterations, and produce the percentile statistic describing where the count lies on the distribution of normal cell alteration counts.

This work presents a viable methodology for identifying cancer cells in scRNA-seq datasets. We identified putative cancer driver alterations in single cells from tumor samples and found that the counts of these alterations differed significantly from what can be found in normal (i.e., non-tumor) cells. Furthermore, there does not appear to be a consistent correlation between inferred CNV and putative driver alteration counts. Our method then provides evidence for the inclusion in downstream analyses of low-inferred-CNV cells that might otherwise be excluded and can strengthen the confidence in calls made on high-inferred-CNV cells.

## Acknowledgments

This work was partly supported by a Cancer Prevention & Research of Texas Recruitment grant (RR200023) to ML and an R37 NIH Cancer Institute grant (5R37CA242070) to ML.

GNU parallel [38] was used for the parallelization of a number of steps described in the methodology presented here.

## Code availability

Code used to generate the primary results presented here is available at github.com/wigasper/calling-cancer.

## Supplementary information

**Fig 6.**
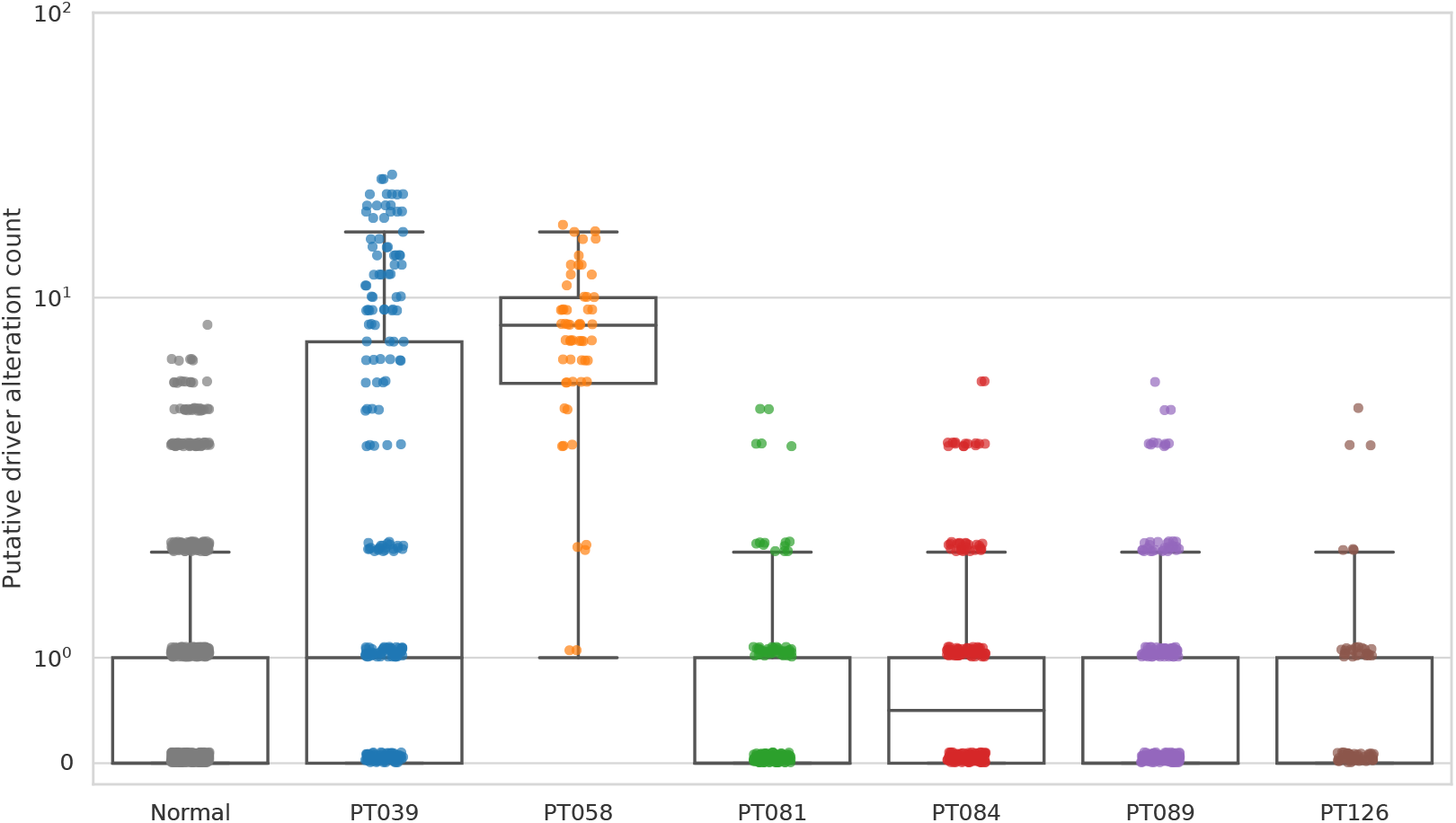
Boxplots showing putative driver alteration counts distributions for all normal tissue cells combined and TNBC dataset cells.

**Fig 7.**
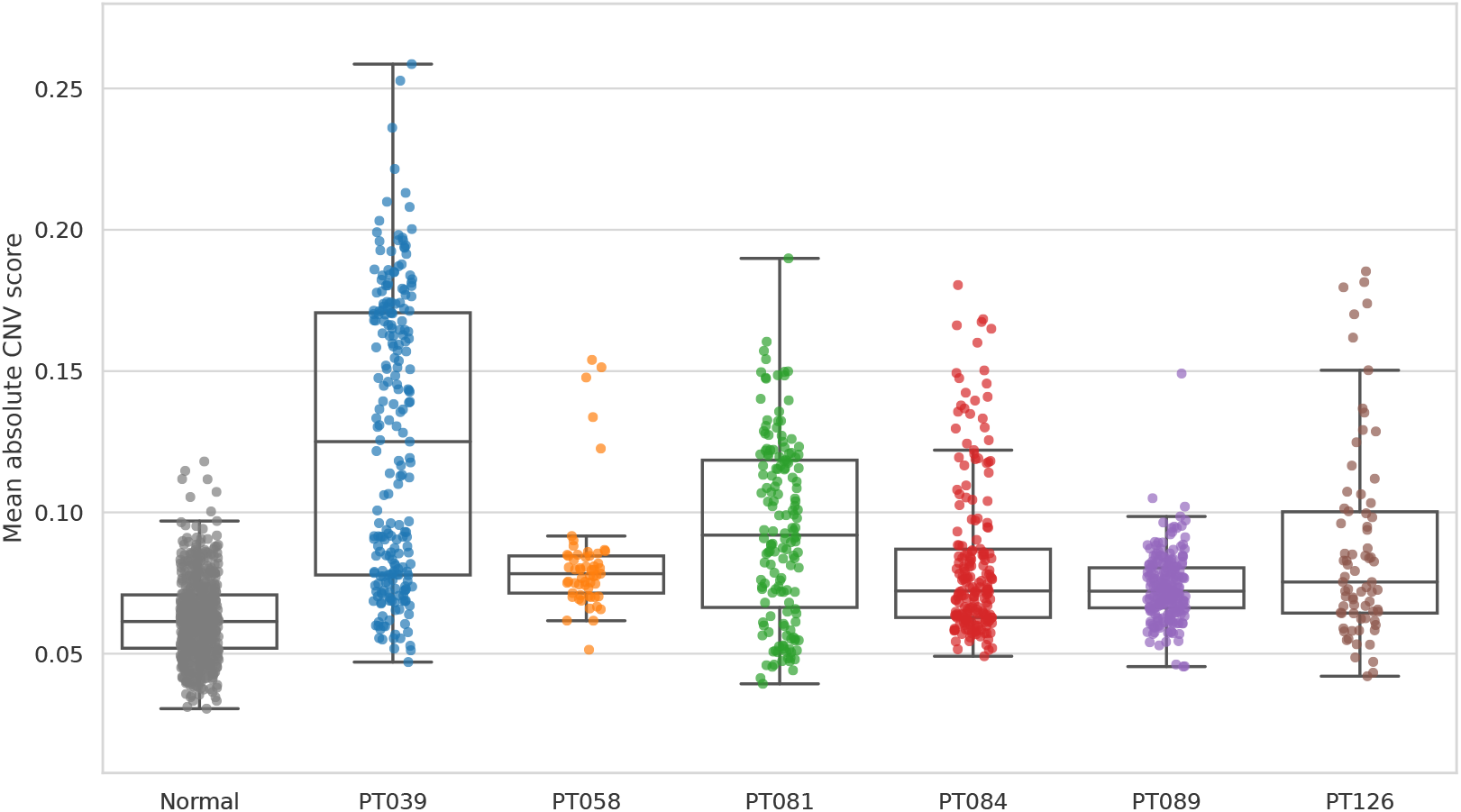
Boxplots showing mean absolute CNV scores for all normal tissue cells combined and TNBC dataset cells.

